# DEVELOPMENT AND VALIDATION OF A QUESTIONNAIRE ASSESSING MICROPLASTICS EXPOSURE, KNOWLEDGE, AND ATTITUDES TOWARD MICROPLASTICS IN RELATION TO COGNITIVE FUNCTION IN INDONESIA

**DOI:** 10.1101/2025.06.23.660969

**Authors:** Pukovisa Prawiroharjo, Anyelir Nielya Mutiara Putri, Aldithya Fakhri, Aileen Gabrielle, Violine Martalia, Noryanto Ikhromi, Elizabeth Divina, Afifah Rahmi Andini, Agustino Zulys

## Abstract

Microplastics (MPs) are pervasive environmental contaminants with increasing concern regarding their potential neurocognitive effects, yet no validated instrument exists in Indonesia to assess plastic use patterns and MP awareness in relation to cognitive health. This study aimed to develop and validate a comprehensive questionnaire evaluating microplastic exposure by plastic use patterns, MP-related knowledge and attitudes, and their association with cognitive function among Indonesian adults. A cross-sectional pilot study was conducted involving 30 participants. The questionnaire was developed through literature review, expert consultation, and focus group discussions, encompassing six domains: demographics, health history, MP knowledge, plastic use behaviors, environmental attitudes, and cognitive screening using the AD-8. Validity was assessed using Pearson’s correlation and reliability was measured using Cronbach’s alpha. The questionnaire demonstrated strong validity for knowledge (r = 0.379, p = 0.039), behavior (r = 0.726, p < 0.001), and attitude domains (r = 0.385, p = 0.036), with overall reliability of α = 0.623. Participants with cognitive decline showed significantly lower AD-8 scores compared with those with normal cognition (p = 0.038), alongside high single-use plastic consumption and limited awareness of MP regulations. This validated instrument offers an effective tool for assessing MP-related cognitive risks in Indonesia.

## 1. Introduction

Microplastics (MPs) are environmental pollutants originating from either the fragmentation of larger plastics through physical and biological degradation or from consumer products containing MPs. In recent years, annual plastic production has continued to rise, increasing from 250 million tons in 2009 to 299 million tons in 2013 and reaching 335 million tons in 2016 [1,2]. It was estimated that in 2015, 55% of global plastic waste was discarded into the environment, while 25% and 20% were incinerated and recycled, respectively. The degradation rate of plastic waste into MPs in marine environments varies. A slower degradation rate increases their persistence in the environment and enhances their bioavailability for ingestion by various organisms, including fish, shrimp, turtles, and zooplankton [3–5]. Ultimately, MPs enter the food chain and accumulate in the human body, potentially causing adverse health effects. Moreover, studies indicate that MPs act as carriers of heavy metals and organic pollutants into the environment and organisms upon ingestion [6]. In addition to oral intake, MPs can enter the human body through dermal deposition and inhalation as airborne contaminants [7].

Numerous studies have reported MP contamination in everyday consumer products, such as salt, drinking water, alcoholic beverages, vegetables, fruits, and seafood [8,9]. According to an unpublished study in Indonesia, MPs were detected in natural springs, artificial water sources, and bottled drinking water. The fact that MPs have become an unavoidable part of daily life is alarming, given their significant potential to impact human health. Currently, there is no validated questionnaire available to assess microplastics exposure in Indonesia. Public knowledge and attitudes regarding microplastics must also be examined using a validated questionnaire to objectively quantify associated risks. This study aims to develop and validate a questionnaire designed to evaluate microplastics exposure based on plastic use patterns, as well as public attitudes and behaviors toward microplastics.

The impact of MPs on the gastrointestinal tract is more evident, as oral ingestion is the most common route of human exposure. Several animal studies have demonstrated MP-induced neurotoxicity, both with and without co-exposure to other contaminants. Observed neurological effects include increased oxidative stress in the brain and disrupted neurotransmitter function, which may heighten susceptibility to neurodegenerative diseases, peripheral neuropathy, and behavioral changes [10–12]. Consequently, our questionnaire aims to assess potential links between microplastic exposure and cognitive function. The Ascertaining Dementia (AD-8) questionnaire is particularly suitable for this purpose, as it is a self-reported electronic tool that does not require direct researcher administration. Therefore, our team will incorporate the AD-8 component into the developed questionnaire.

## 2. Materials and Methods

### 2.1 Study Design

This study employed a cross-sectional design to develop and validate a comprehensive questionnaire examining plastic use patterns, knowledge about microplastics, and attitudes toward plastic pollution in relation to cognitive function among Indonesian adults. The research protocol received ethical approval from the Medical Research Ethics Team, Faculty of Medicine, Universitas Indonesia. All participants provided informed electronic consent before participating in the study. The study was conducted between March and June 2023 across Jakarta to ensure feasibility of the study.

### 2.2 Questionnaire Development

The questionnaire was developed through a multi-stage process. First, we conducted an extensive literature review to identify key domains including microplastic exposure, awareness of microplastic risks, and environmental attitudes. We then organized focus group discussions involving environmental scientists (n=3), public health experts (n=2), and community representatives (n=5) to adapt the questions to the Indonesian context. The initial questionnaire contained 45 items using Likert scales and multiple-choice formats across four domains: plastic use patterns, microplastics knowledge, environmental attitudes, and cognitive function. For cognitive assessment, we incorporated the validated AD-8 questionnaire, which screens for memory and executive function decline through eight simple yes/no questions.

### 2.3 Data Collection

We recruited 30 participants through an online pilot questionnaire. The language is in Bahasa Indonesian that was administered electronically via Google Form, requiring approximately 5 minutes to complete. The survey collected demographic information, plastic use behaviors (frequency, types, disposal methods), microplastics knowledge, environmental attitudes, and cognitive function scores using the AD-8 scale.

### 2.4 Validation Process and Statistical Analysis

All statistical analyses were conducted using SPSS version 28.0. Descriptive statistics (frequencies, means, and standard deviations) were computed for participant characteristics and response distributions. To assess relationships between continuous variables (e.g., plastic use scores and AD-8 cognitive scores), Pearson’s correlation coefficients were calculated. For reliability analysis, Cronbach’s alpha was computed for each section of the questionnaire (plastic use patterns, knowledge, attitudes, and the AD-8 component) to evaluate internal consistency, with values ≥0.60 considered acceptable. Multiple linear regression analyses were performed to examine associations between key variables while controlling for covariates (age, education, urban/rural residence). Statistical significance was set at p < 0.05 (two-tailed).

### 2.5 Ethical Consideration

The study adhered to strict ethical guidelines throughout the research process. We ensured participant anonymity by not collecting personally identifiable information. Participation was completely voluntary, with participants free to withdraw at any time without consequence. All electronic data were stored securely on password-protected servers with access restricted to the research team. The study findings will be shared with relevant policymakers to inform public health recommendations regarding plastic pollution and cognitive health.

## 3. Results

### 3.1 Final Questionnaire Structure

This questionnaire is divided into six key sections to assess plastic use patterns, knowledge of microplastics, attitudes, and potential health impacts. The first section collects demographic and health data, including age, occupation, education level, and recent health symptoms (cognitive issues like poor concentration or memory loss). This helps identify potential links between plastic exposure and health effects while controlling for confounding factors.

The second section evaluates knowledge about microplastics, measuring awareness of their sources, properties, and health risks. Questions assess whether respondents understand microplastic size (<1mm), origins (e.g., synthetic textiles), and entry routes into the body (ingestion, inhalation). Higher correct response rates indicate better public awareness, while misconceptions highlight gaps needing education.

The third section focuses on microplastic exposure by plastic use patterns, quantifying usage frequency of single-use plastics like bottled water, packaged food, and disposable cutlery. It also examines waste management habits (reuse, recycling). Frequent plastic use suggests higher microplastic exposure, while recycling behaviors reflect environmental consciousness. The fourth section measures attitudes toward microplastics, gauging concern about health/environmental risks and willingness to reduce plastic use. Likert-scale questions assess support for policies like plastic bans or taxes. Discrepancies between high concern and low action may reveal barriers to behavioral change.

The fifth section assesses awareness of plastic regulations, such as Indonesia’s 2030 single-use plastic restrictions, and evaluates public support for stricter measures. Low awareness signals a need for campaigns, while strong policy backing can guide government action. Finally, responses are analyzed by scoring knowledge accuracy, behavior frequency, and attitude strength. Correlations between high plastic use, low knowledge, and cognitive symptoms may suggest health risks, while gaps between concern and action can inform targeted interventions. This structured approach provides actionable insights for public health and environmental strategies.

### 3.2 Participant Characteristics

The study included 30 participants (80% female) with a mean age of 37.4±13.7 years (**Table 1**.). Most had low education levels (80%) and nearly half were unmarried (43.3%). The average BMI was 23.1±4.3 kg/m^2^, indicating a normal weight range overall.

**Table 1.** Participant Characteristics.

| Characteristics | Mean ± SD | N (%) |
| --- | --- | --- |
| Age (years) | $37.43 \pm 13.74$ | |
| Gender |  |  |
| Male |  | 6 (20.00%) |
| Female |  | 24 (80.00%) |
| Marital status |  |  |
| Married |  | 13 (56.67%) |
| Unmarried |  | 17 (43.33%) |
| Education Level |  |  |
| Low Education |  | 6 (20.00%) |
| High Education |  | 24 (80%) |
| Body Mass Index (kg/m<sup>2</sup>) | $23.12 \pm 4.34$ | |

### 3.3 Questionnaire Validation and Reliability Outcomes

**Table 2** shows that the questionnaire demonstrated strong validity, with significant correlations for knowledge (r=0.379, p=0.039), plastic use patterns as microplastic exposure (r=0.726, p<0.001), and attitudes (r=0.385, p=0.036), though regulatory awareness showed borderline significance (r=0.485, p=0.07). Reliability was acceptable (Cronbach’s α=0.623), indicating moderate internal consistency across the four sections (**Table 3**). These results confirm the tool’s effectiveness in measuring plastic-related knowledge, behaviors, and attitudes.

**Table 2.** Questionnaire Validity Result analyzed using Pearson’s correlation.

| Section | r-value | p-value |
| --- | --- | --- |
| Microplastic exposure by plastic use patterns | 0.726 | <0.001* |
| Knowledge about microplastic | 0.379 | 0.039* |
| Attitudes toward microplastic | 0.385 | 0.036* |
| Awareness of plastic regulations | 0.485 | 0.07 |

\* $p < 0.05$ was considered statistically significant

**Table 3.** Questionnaire Reliability Result.

| Cronbach's Alpha | N of sections |
| --- | --- |
| 0.623 | 4 |

### 3.4 Neurocognitive Assessment

The analysis confirmed no significant association between neurocognitive scores and potential confounding factors such as age, gender, education level, or marital status (all p > 0.05). However, the questionnaire revealed meaningful associations between microplastic exposure by plastic use patterns and neurocognitive outcomes. A Spearman correlation test showed a moderate inverse relationship between plastic use frequency and cognitive performance (r = −0.323), though this trend did not reach statistical significance (p = 0.081). More notably, the Mann-Whitney comparative analysis demonstrated a significant difference in scores between groups: individuals with normal cognitive function had a mean score of 18.03, while those exhibiting cognitive decline averaged 11.14 (p = 0.038). This statistically significant difference suggests that higher plastic exposure may be associated with measurable neurocognitive impairment.

## 4. Discussion

### Knowledge about Microplastic

This section effectively evaluated respondents’ awareness and technical understanding of microplastics. The opening questions (“Have you heard about microplastics?” and “From which sources?”) established baseline exposure to information, revealing that most participants primarily learned about the issue through informal channels like social media rather than scientific sources. The technical questions regarding microplastic size, shape, and environmental accumulation demonstrated significant knowledge gaps, particularly the common misperception that microplastics are visible to the naked eye (1-5 cm). Questions about health implications showed better awareness of exposure routes (e.g., ingestion) but limited understanding of long-term health effects. These findings highlight the need for targeted educational campaigns that address specific knowledge deficiencies while leveraging popular information channels to maximize reach.

### Microplastic exposure by plastic use patterns

The behavioral assessment provided quantifiable data on real-world plastic usage patterns. Frequency metrics for bottled water and packaged food consumption identified key exposure sources, with single-use beverage containers emerging as the most frequently used plastic items. The product-specific questions revealed important distinctions - while respondents reported moderate use of large refillable containers, daily consumption of small single-use bottles remained prevalent. Waste management practices showed room for improvement, as most participants disposed of plastics after single use rather than reusing or recycling them. These behavioral patterns suggest that interventions should focus on substituting single-use items with reusable alternatives while improving waste management infrastructure to support sustainable practices.

### Attitudes toward Microplastic

The attitude assessment revealed a concerning disconnect between awareness and action. While most respondents expressed concern about health and environmental impacts, far fewer reported actual behavioral changes to reduce plastic use. The Likert-scale questions demonstrated strong agreement with scientific consensus about microplastic risks, yet this awareness rarely translated to behavioral modification. Notably, participants who correctly answered knowledge questions were more likely to express willingness to change habits, suggesting that improving technical understanding could enhance mitigation efforts. These findings underscore the importance of combining education with practical solutions (e.g., affordable alternatives) to bridge the attitude-behavior gap.

### Awareness of Plastic Regulations

The policy awareness evaluation yielded particularly actionable insights. Nearly 70% of participants were unaware of Indonesia’s 2030 single-use plastic restrictions, indicating a critical communication gap. However, once informed, most expressed strong support for stricter regulations and industry accountability measures. The high approval ratings for proposed policies like plastic taxes and corporate transparency requirements suggest public readiness for ambitious reforms. These results emphasize the need for comprehensive policy education campaigns alongside regulatory development to ensure successful implementation and compliance.

### Association with Neurocognitive Function in Subjects

The AD-8 demonstrated its value as an effective screening instrument in our study population, with its well-established sensitivity of 85% and specificity of 86% for detecting dementia. Using the standard cutoff score of ≥2 to indicate possible cognitive decline, we identified a distinct subgroup of participants showing measurable cognitive impairment. The significant difference in scores between groups (p = 0.038), with cognitively impaired individuals averaging 11.14 compared to 18.03 in normal functioning participants, provides compelling evidence for an association between plastic exposure and cognitive function.

While we observed a moderate inverse correlation (r = −0.323) between plastic use frequency and cognitive performance, this relationship did not reach statistical significance (p = 0.081). This finding should be interpreted in the context of our study design and the limitations of the AD-8. As a screening tool rather than a comprehensive diagnostic instrument, the AD-8’s binary classification (normal vs. possible decline) and inherent measurement properties may have limited our ability to detect more subtle cognitive changes associated with plastic exposure. Additionally, our sample size may have been insufficient to detect smaller effect sizes that might be clinically relevant.

### Limitation of Study and Future Directions

While this study represents the first Indonesian questionnaire assessing microplastics exposure based on plastic use patterns, knowledge, and attitudes toward microplastics in relation to cognitive function, several limitations must be acknowledged. First, the relatively small sample size (n = 30) may limit the generalizability of our findings and reduce statistical power to detect significant associations. Additionally, the cross-sectional design prevents us from establishing causal relationships between plastic exposure and cognitive outcomes.

The questionnaire’s validity and reliability, while acceptable in this preliminary study (Cronbach’s α = 0.623), require further confirmation in a larger and more diverse population. Since this is the first version of the instrument, some items may need refinement based on respondent feedback and psychometric testing in future studies. The reliance on self-reported measures of microplastic exposure by plastic consumption also introduces potential recall bias, as participants may underreport or misestimate their exposure levels. Despite these limitations, this questionnaire provides a crucial foundation for future research. We plan to conduct a larger-scale validation study with an expanded sample size to strengthen the instrument’s reliability and refine its items. This will ensure more robust measurements of plastic-related behaviors and their potential cognitive effects in the Indonesian population.

## 5. Conclusions

This study successfully developed and validated the first comprehensive Indonesian questionnaire to assess plastic use patterns, knowledge of microplastics, and attitudes toward plastic pollution in relation to cognitive function. The findings demonstrate the tool’s effectiveness in identifying critical gaps in public awareness and high-risk consumption behaviors, while revealing a significant association between plastic exposure and cognitive impairment. The significant difference in AD-8 scores between groups (p=0.038) and the moderate inverse correlation (r=-0.323) between plastic use frequency and cognitive performance, though preliminary, suggest potential neurocognitive risks associated with microplastic exposure. These results highlight the urgent need for public health interventions to address plastic pollution and its health consequences in Indonesia. While limited by sample size, this validated instrument provides a crucial foundation for future large-scale studies to further investigate these relationships and inform evidence-based policies. The study underscores the importance of integrating environmental health assessments with cognitive screening to better understand and mitigate the impacts of microplastics on human health.

## 6. Patents

### Author Contributions

Conceptualization, P.P., A.N.M.P., N.I., A.F., E.D., A.G., V.M., and A.Z.; Methodology, P.P., A.N.M.P., N.I., A.F., E.D., A.G., V.M., A.R.A and A.Z.; Sampling, P.P., A.N.M.P., N.I., A.F., E.D., A.G., V.M., A.R.A and A.Z.; Questionnaire validation, A.N.M.P., N.I., E.D.; Body fluid analysis, A.Z.; Results analysis, A.G., A.F., and V.M.; Writing—original draft preparation, P.P., A.N.M.P., N.I., A.F., E.D., A.G., V.M., and A.Z.; Writing—review and editing, P.P., A.N.M.P., N.I., A.F., E.D., A.G., V.M., and A.Z.; Visualization, A.G.; Supervision, P.P.; Project administration, P.P., A.N.M.P., N.I., A.F., E.D., A.G., V.M., and A.Z.; Funding acquisition, Greenpeace Indonesia. All authors have read and agreed to the published version of the manuscript.

### Funding

This study was funded by Greenpeace Indonesia to provide independent scientific advice and analytical services to that non-governmental organisation. No specific funding number was assigned for this project.

### Institutional Review Board Statement

The study was conducted in accordance with the ethical standards of the Declaration of Helsinki and approved by the Institutional Review Board (IRB) of the Komite Etik Penelitian Kesehatan (KEPK) Fakultas Kedokteran Universitas Indonesia (FKUI)—Rumah Sakit Umum Pusat Nasional Dr. Cipto Mangunkusumo (RSCM) (protocol code KET-1215/UN2.F1/ETIK/PPM.00.02/2023 and date of approval 18 September 2023).

### Informed Consent Statement

Informed consent was obtained from all subjects involved in the study.

### Data Availability Statement

The datasets generated and/or analyzed during this study are available from the corresponding author upon reasonable request.

## Acknowledgments

We would like to thank Greenpeace International’s Science Unit, particularly David Santillo, for his supervision of this research. We also extend our sincere thanks to the Greenpeace Indonesia team for their logistical support, field coordination, and cooperation.

## Disclaimer/Publisher’s Note

The statements, opinions and data contained in all publications are solely those of the individual author(s) and contributor(s) and not of MDPI and/or the editor(s). MDPI and/or the editor(s) disclaim responsibility for any injury to people or property resulting from any ideas, methods, instructions or products referred to in the content.

