## Supplementary Material for "DEVELOPMENT AND VALIDATION OF A QUESTIONNAIRE ASSESSING MICROPLASTICS EXPOSURE, KNOWLEDGE, AND ATTITUDES TOWARD MICROPLASTICS IN RELATION TO COGNITIVE FUNCTION IN INDONESIA"

**Questionnaire on Knowledge, Behavior, and Attitudes Related to Microplastics**

**Part 1.**

**Identity**

Name	:
Age	:
Gender	: Male / Female
Marital Status	: Married / Unmarried
Occupation	:
Body Weight	:
Height	:
Education Level: Elementary / Junior High / Senior High / Diploma / Bachelor / Master / Doctoral
Address	:
Mobile Number:

**Medical History**

1. **Within the past 1 month, have you experienced any of the following complaints? (You may check more than one)**
   ☐ Nausea and vomiting
   ☐ Diarrhea
   ☐ Constipation
   ☐ Abdominal bloating
   ☐ Urinary problems
   ☐ Sleep disturbances
   ☐ Decreased concentration
   ☐ Blurred vision
   ☐ Hearing problems
   ☐ Headache
   ☐ Vertigo (spinning dizziness)
   ☐ Frequent forgetfulness
   ☐ Disorientation / confusion
   ☐ Easily fatigued during activities
   ☐ Seizures
   ☐ Recurrent fainting
   ☐ Menstrual disorders
   ☐ Not yet / difficulty having children (for those who are married)
2. **Do you have a history of illness that currently requires routine treatment?**
   (e.g., epilepsy, stroke, head injury, heart disease, chronic infection, etc.): *open-ended*
3. **Do you have a history of congenital (birth-related) disease?** *open-ended*

**Part 2. Plastic Use Patterns (behavior related to the use of plastic packaging)**

| Types of Plastics based on the plastic resin identification code (RIC) system | More than 3 times/day | 1–3 times/day | 4–6 times/week | 1–3 times/week | 1–3 times/month | <1 time/month | never |
| --- | --- | --- | --- | --- | --- | --- | --- |
| [1] Packaged beverages and in plastic (commonly sold by street vendors), mineral water, fruit juice containers, and cooking oil  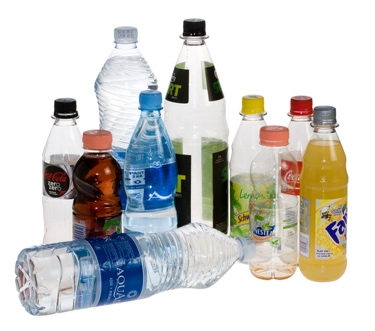  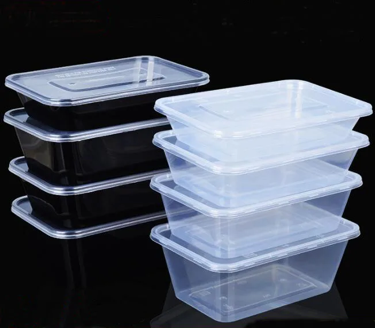 |  |  |  |  |  |  |  |
| [2] Milk jugs, cleaning agents, laundry detergents, bleaching agents, shampoo bottles, washing soaps  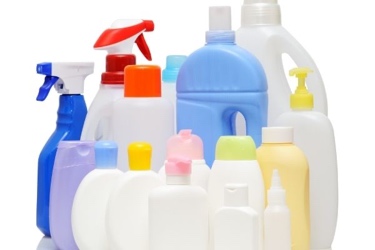 |  |  |  |  |  |  |  |
| [3] Trays for sweets, fruit, plastic packing (bubble foil) and food foils to wrap food stuff  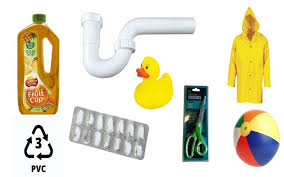 |  |  |  |  |  |  |  |
| [4] Crushed bottles, shopping bags, highly-resistant sacks and most of the wrappings  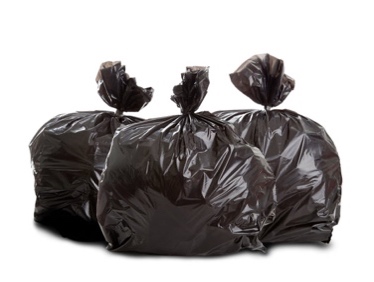  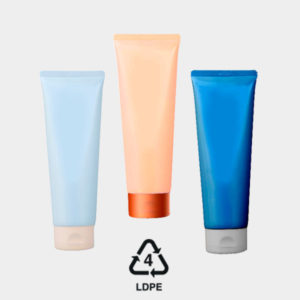  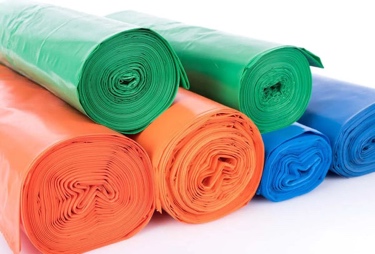 |  |  |  |  |  |  |  |
| [5] Heat-resistant food containers, yogurt cups, medicine bottles, and reusable plasticware.  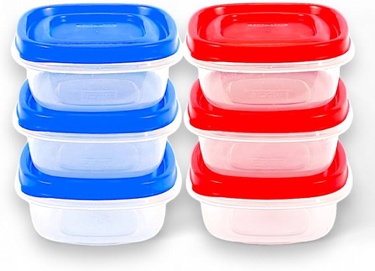  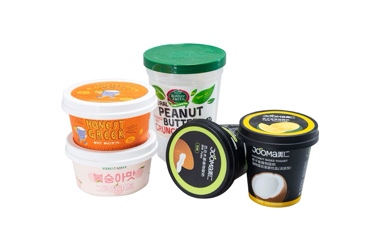 |  |  |  |  |  |  |  |
| [6] Disposable Styrofoam food boxes, plastic cutlery, egg trays, and single-use coffee cups.  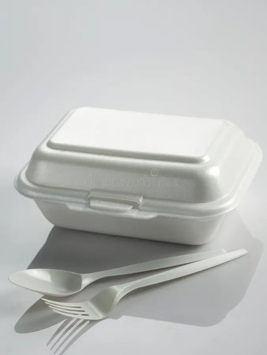  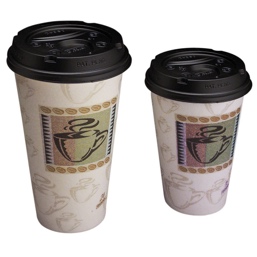 |  |  |  |  |  |  |  |
| [7] Large reusable water jugs, sports bottles, and specialized polycarbonate products.  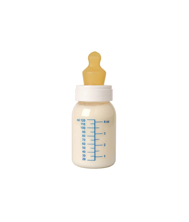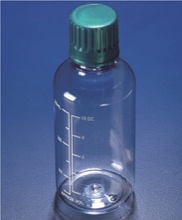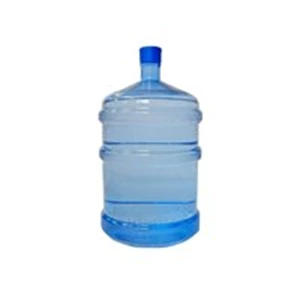 |  |  |  |  |  |  |  |

**Part 3. Knowledge Related to Microplastics**

1. **Have you ever heard of microplastics?**
   A) Yes, often
   B) Yes, occasionally
   C) No
2. **From which sources have you heard about microplastics?**
   A) Friends and relatives
   B) Conferences and seminars
   C) Internet
   D) Books
   E) Informational brochures
   F) Radio
   G) Television
   H) Newspapers
   I) Environmental journals
   J) Others: ……
3. **Have you ever searched for information about microplastics?**
   A) Always
   B) Sometimes (>50%)
   C) Rarely (<50%)
   D) Never
4. **In your opinion, what forms do microplastics have?**
   A) Fibers

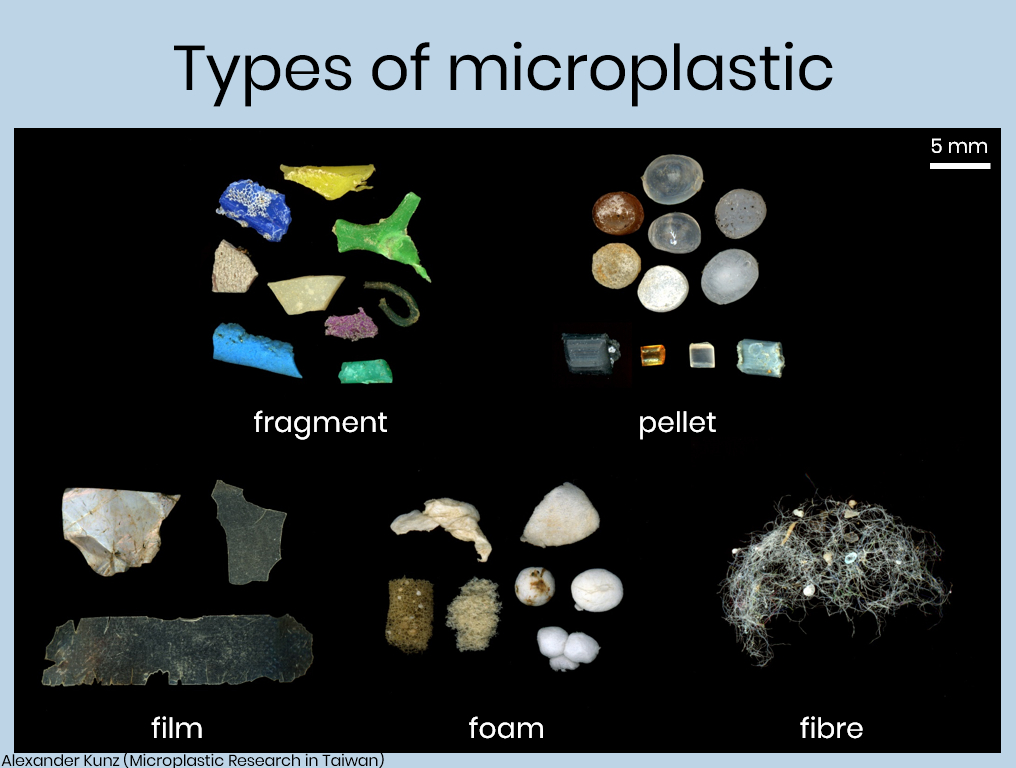

B) Spherical particles

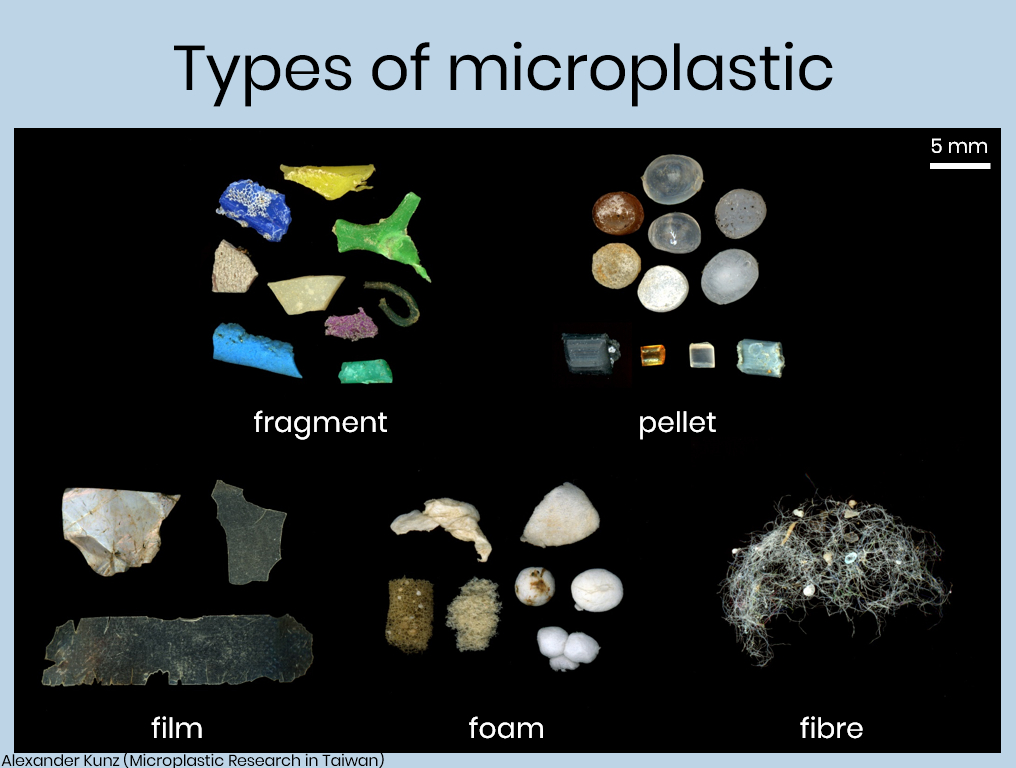

C) Irregular particles

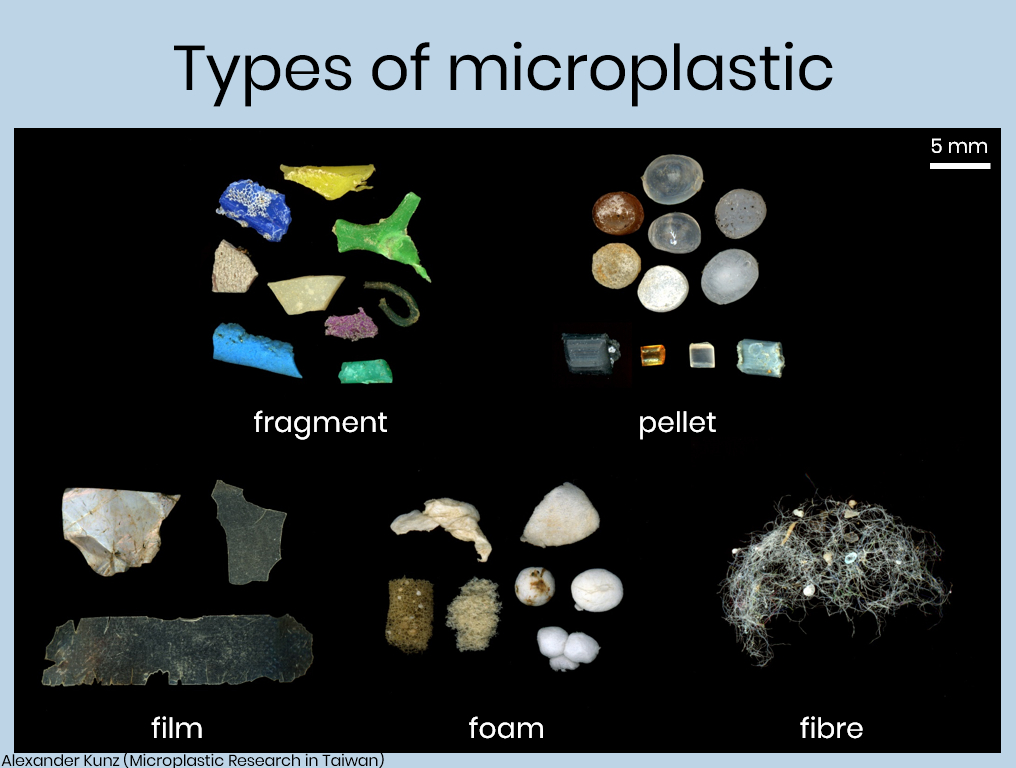

D) Films

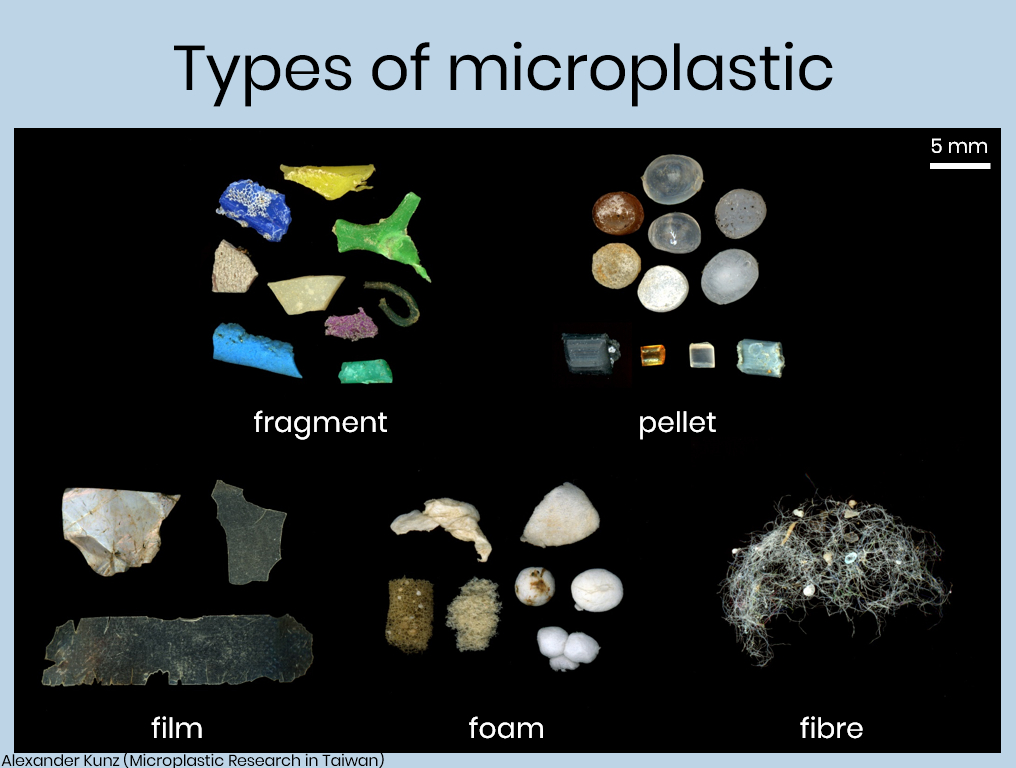

5. **In your opinion, what is the size of microplastics?**
   A) Less than 1 mm
   B) Around 1–5 cm
   C) More than 10 cm
6. **In your opinion, what are the sources of microplastics? (You may choose more than one)**
   A) Industrial plastic production
   B) Release of fibers and particles into water
   C) Washing of synthetic or plastic-blended clothing such as polyester
   D) Use of plastic-based products
   E) Use of personal care products; microbeads or fine scrub particles
7. **In your opinion, where can microplastics accumulate in the environment?**
   A) Freshwater
   B) Seawater
   C) Air
   D) Soil
   E) Marine biota
   F) Salt
   G) Animal products such as meat, eggs, and milk
8. **Can microplastics affect human health?**
   A) Yes
   B) No
   C) Do not know
9. **Are you aware that microplastics have been found in the human body?**
   A) Yes
   B) No
10. **In your opinion, how can microplastics enter the human body?** (You may select more than one answer)
    A) Mouth – gastrointestinal tract
    B) Nose – respiratory tract
    C) Skin

**Criteria Used in Choosing Plastic Packaging**

| Criteria | Strongly agree | Agree | Neutral | Disagree | Strongly disagree |
| --- | --- | --- | --- | --- | --- |
| Price |  |  |  |  |  |
| Brand |  |  |  |  |  |
| Quality |  |  |  |  |  |
| Availability/ easy access |  |  |  |  |  |
| Personal preference |  |  |  |  |  |
| Visuals |  |  |  |  |  |

1. **Where do you get information regarding the quality of plastic-packaged drinking water? (You may choose more than one)**
   A) Television
   B) Books / journals
   C) Newspapers
   D) Internet
   E) Labels on packaging bottles
   F) Friends / relatives
   G) Others ………
2. **How do you manage your drinking water packaging waste?**
   A) Refilled
   B) Reused for other purposes (plant pots, crafts, etc.)
   C) Disposed of after use

**Part 4. Attitudes Toward Microplastics**

1. **Are you concerned about the negative environmental impacts of using plastic-packaged drinking water?**
   A) Yes
   B) No
   C) Do not know
2. **Are you concerned about the negative health impacts of using plastic-packaged drinking water?**
   A) Yes
   B) No
   C) Do not know
3. **How often do you manage waste from plastic-packaged drinking water?**
   A) Always
   B) Sometimes (>50%)
   C) Rarely (<50%)
   D) Never
4. **After knowing that microplastics have been found in daily consumer products, what is your attitude?**
   A) Reduce plastic use
   B) Stop consuming products indicated to contain microplastics
   C) Continue using/consuming these products

**Statements:**

| Statement: | Strongly agree | Agree | Neutral | Disagree | Strongly disagree | Do not know |
| --- | --- | --- | --- | --- | --- | --- |
| **Among all drinking water packaging, I most frequently use refillable gallon packaging** |  |  |  |  |  |  |
| Microplastics can contain and accumulate hazardous chemicals |  |  |  |  |  |  |
| Microplastics can absorb and transport other contaminants |  |  |  |  |  |  |
| Microplastics do not degrade in the environment |  |  |  |  |  |  |
| Microplastics endanger human health through consumption and inhalation |  |  |  |  |  |  |
| Microplastics endanger animal health |  |  |  |  |  |  |
| Microplastics harm the economy |  |  |  |  |  |  |

**Part 5. Knowledge and Attitudes Toward Plastic Management Regulations in Indonesia**

1. **Are you aware that by 2030 there will be regulations restricting certain types of single-use plastics in Indonesia?**
   A) Yes
   B) No
2. **Which types of single-use plastics do you know are banned/restricted?**
   A) Plastic bags
   B) Styrofoam
   C) Plastic packaging under 50 ml (sachets)
   D) Multilayer plastic packaging
   E) Beverage bottles under 500 ml
   F) Do not know
3. **Do you agree with the regulation restricting single-use plastics?**
   A) Yes
   B) No
4. **In your opinion, should the regulation restricting single-use plastics be expanded nationwide across Indonesia?**
   A) Yes
   B) No
5. **Given the current plastic pollution crisis, do you think restrictions on single-use plastics should be accelerated to before 2030?**
   A) Yes
   B) No

**Do you agree with the following policy options to reduce microplastic pollution?**

| **Policy Option** | **Strongly Agree** | **Agree** | **Neutral** | **Disagree** | **Strongly Disagree** | **Do Not Know** |
| --- | --- | --- | --- | --- | --- | --- |
| **Limiting the use of single-use plastic packaging** |  |  |  |  |  |  |
| **Stopping the production of virgin plastic by 2030** |  |  |  |  |  |  |
| **Encouraging industries to provide refill/reuse systems** |  |  |  |  |  |  |
| **Imposing excise taxes on industries that still rely on plastic** |  |  |  |  |  |  |
| **Establishing standards for microplastic contamination in food** |  |  |  |  |  |  |
| **Establishing standards for microplastic contamination in water bodies/environment** |  |  |  |  |  |  |
| **Improving water purification technology to prevent microplastic contamination in drinking water sources** |  |  |  |  |  |  |
| **Requiring industry transparency regarding chemical content and risks of plastic packaging, including potential microplastic pollution** |  |  |  |  |  |  |
| **Improving local waste collection, sorting, and management systems** |  |  |  |  |  |  |

**Part 6. cognitive screening using the AD-8.**

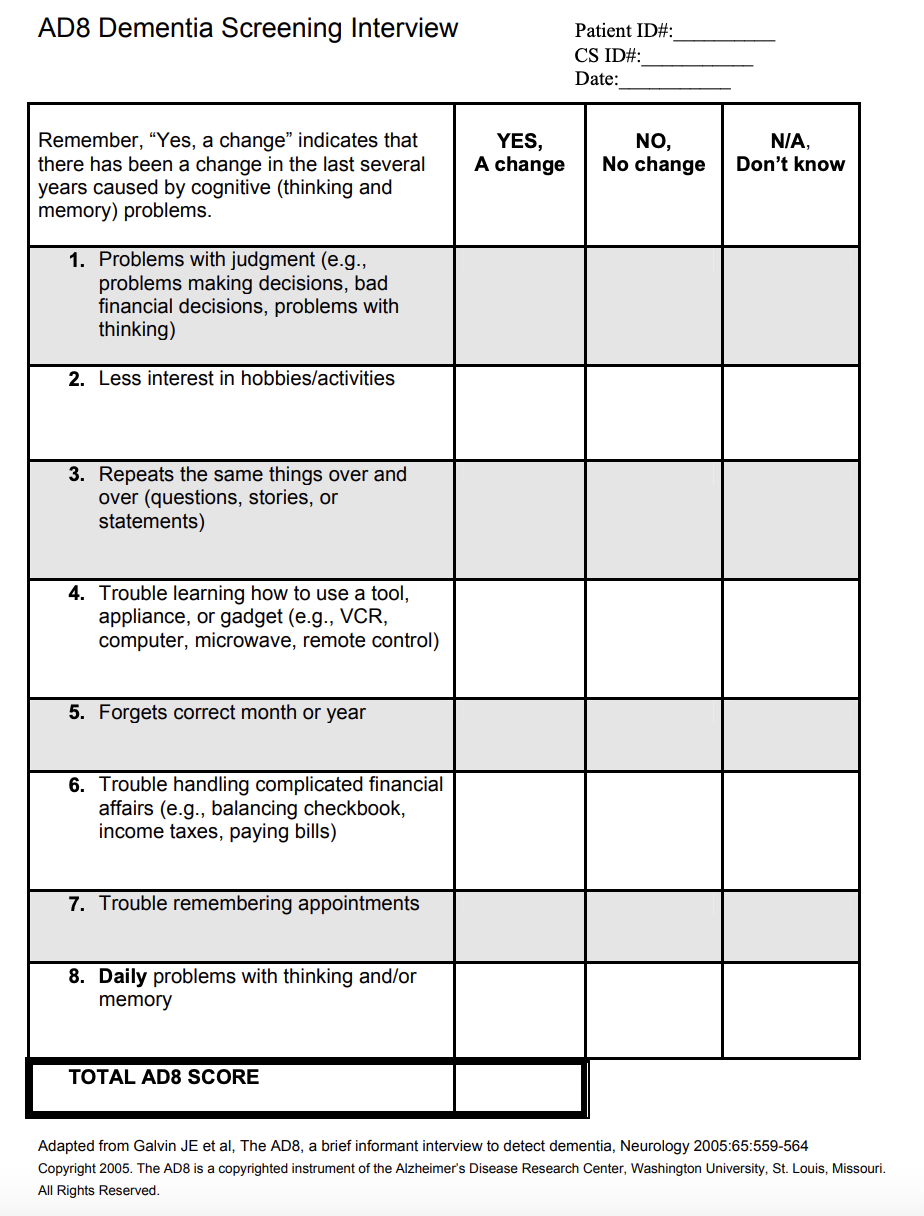

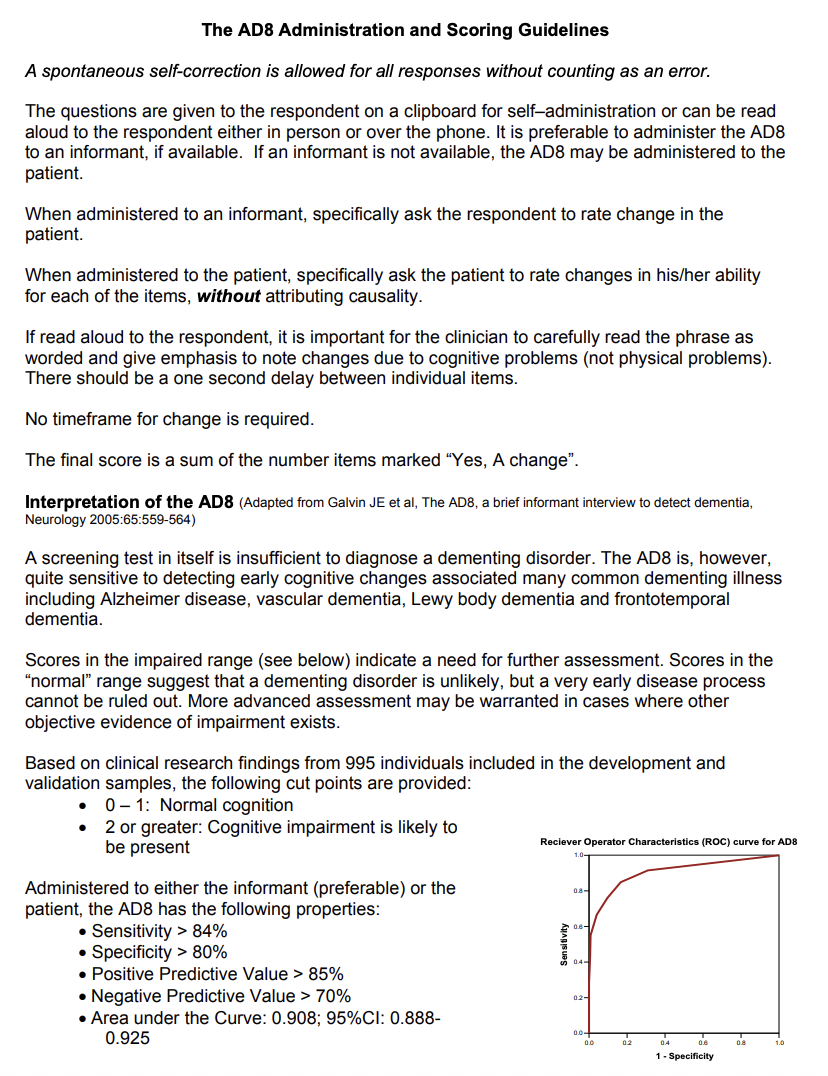

**Microplastic Exposure Assessment and Scoring**

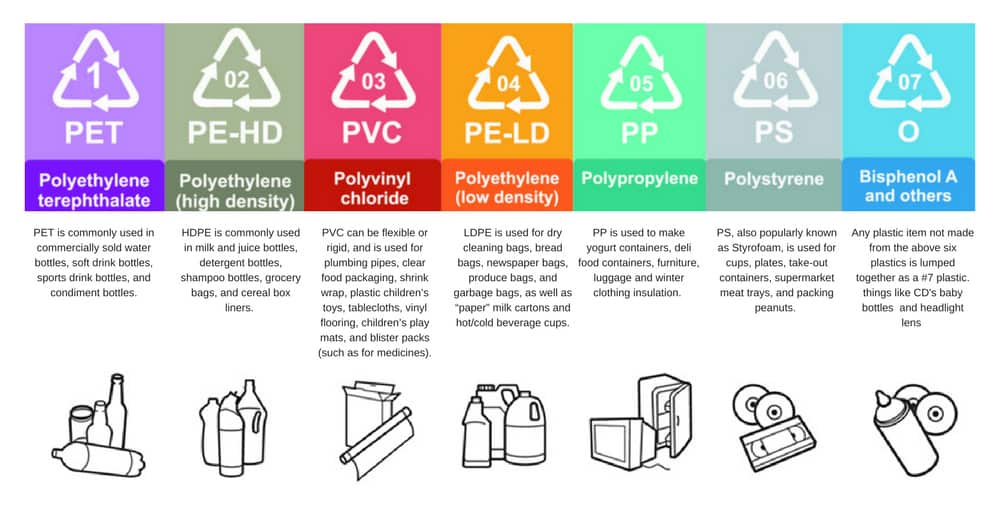
**Part 2: Behavior Related to the Use of Plastic-Packaged Drinking Water.**

**Figure 1. Classification of plastic resin codes.**

**Participants reported their consumption of seven categories of plastic products, classified according to the plastic resin identification code (RIC) system.**

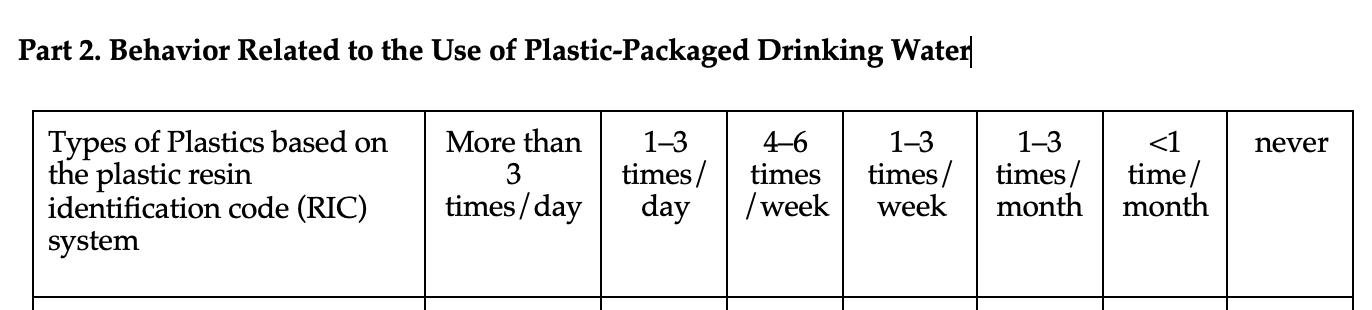

**Microplastic exposure was scored based on self-reported frequency of use and categorized as follows:**

- Never, <1 time per month, or 1–3 times per month: Low exposure (score = 1)
- 1–3 times per week or 4–6 times per week: Moderate exposure (score = 2)
- 1–3 times per day or >3 times per day: High exposure (score = 3)

The frequency-based exposure scores across all plastic product categories were summed to generate a composite microplastic exposure score. Accordingly, the total microplastic exposure score ranged from 7 (minimum) to 21 (maximum), with higher scores indicating greater potential exposure.
